# Optimising the use of Oxford nanopore sequencing technology for detection of *Wolbachia* bacterial endosymbionts in *Anopheles* mosquitoes

**DOI:** 10.1101/2025.06.16.659891

**Authors:** Laura Chatterley, Nicola Hull, Isabel Hughes, Seynabou Sougoufara, Owain Meek, Murat Ceyran, Vishaal Dhokiya, Janvier Bandibabone, Cyprian Adala, Elin Cunningham, Shehu Shagari Awandu, Antonio Nkondjio, Grant Hughes, Eva Heinz, Thomas Walker

## Abstract

Malaria, a mosquito-borne disease predominantly affecting sub-Saharan Africa, remains a significant global health challenge with 263 million cases in 2023. Although progress in vaccine development has recently occurred, insecticide resistance has reduced the effectiveness of the main stay method of traditional vector control and novel innovative vector control strategies against *Anopheles* mosquitoes is crucial to combat the spread of malaria. *Wolbachia* is an endosymbiotic bacterium that can invade mosquito populations and inhibit human pathogens. Historically *Wolbachia* was thought to not occur naturally within wild *Anopheles An*.) populations, but recent evidence has confirmed the presence of at least two strains in genuine symbiosis. Detection of *Wolbachia* strains has pre-dominantly relied on PCR amplification of *Wolbachia* genes and/or sanger and illumina sequencing. In this study, we assess the use of Oxford Nanopore Technology (ONT) for detecting bacteria within wild *Anopheles* mosquitoes and determine the ability to detect the endosymbiotc bacterium *Wolbachia*. We assessed *Wolbachia* detection from library preparations using the ONT Field sequencing kit for MinION sequencing on the Mk1C and found a pool size of 10 mosquitoes is required preventing individual mosquito analysis. There were roughly 13 times more *Wolbachia* reads in pooled *Anopheles demeilloni* compared to *An. gambiae*. Utilizing the more economical Flongle flow cells and Mk1B device, we further evaluated the effectiveness of two library preparation methods that are compatible: Rapid PCR Barcoding and 16S Barcoding kits. Our findings indicate that the 16S Barcoding Kit in conjunction with the Flongle flowcell effectively identified *Wolbachia* in both *Anopheles demeilloni* and *Anopheles moucheti* but we found no evidence of *Wolbachia* in *Anopheles gambiae*. Our study demonstrates the feasibility of developing ONT for field-based sequencing in malaria-endemic regions, providing a potentially lower cost and ‘field friendly’ method for monitoring mosquito populations and associated endosymbionts.

**Graphical abstract:** 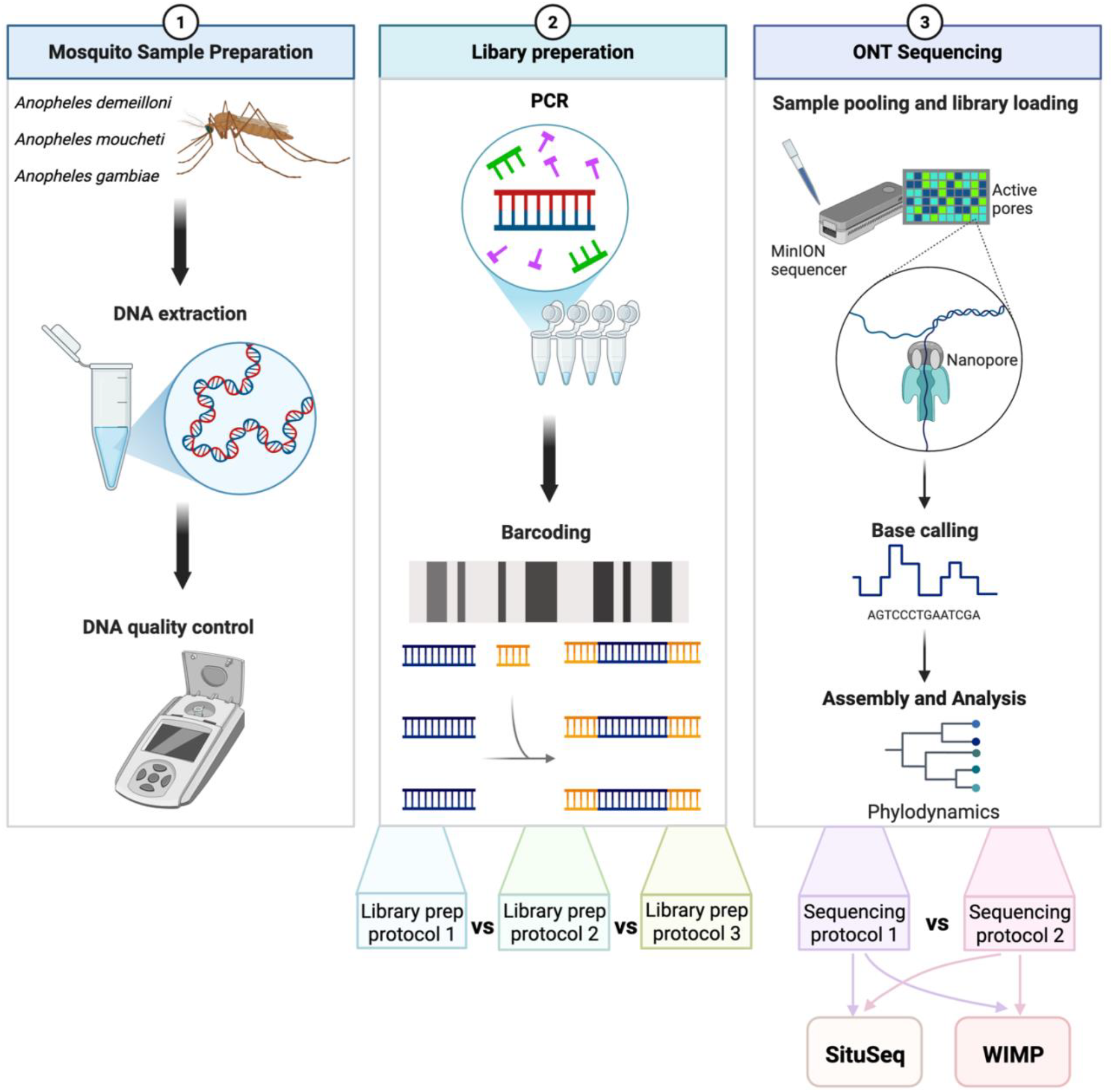

## Introduction

Malaria is caused by *Plasmodium* parasites transmitted through the bites of infected *Anopheles (An*.) mosquitoes and there were an estimated 263 million malaria cases and 597,000 deaths globally in 2023^1^. Despite encouraging progress in vaccine development, insecticide resistance has reduced the effectiveness of traditional vector control. Bacteria that naturally reside within *Anopheles* mosquitoes have been shown to inhibit *Plasmodium* but the major hurdle has been the lack of a natural mechanism to spread bacteria through mosquito populations. *Wolbachia* endosymbiotic bacteria can invade mosquito populations through phenotypic adaptations such as cytoplasmic incompatibility (CI), which results from the sterility of progeny from matings between *Wolbachia*-infected males and uninfected females (all other crosses are compatible) providing a natural drive mechanism. In *Aedes aegypti* mosquitoes that have been artificially transinfected, *Wolbachia* both invade wild populations and inhibit arbovirus transmission^2-6^. Integrated *Anopheles* vector management requires the development of novel control strategies that can work synergistically with existing methods. *Wolbachia*-based biocontrol is safe for humans and already widely accepted. Importantly, it also does not require complex, logistically challenging and highly infrastructure-dependent distribution campaigns or specific assessments to target specific population groups. For replacement strategies only small release numbers are needed making this strategy sustainable, eco-friendly and without bias against specific population groups or hard-to-reach settlements. Based on *Wolbachia*-infected *Aedes* release programmes, it is feasible that a single release can be enough to trigger population invasion over many kilometres and the scalability is much larger than other control methods making this economically cost effective. Furthermore, dengue suppression via *Wolbachia*-infected *Ae. aegypti* release is the only method that has been shown to effectively reduce incidence across multiple countries giving great hope that this could be effective against *Anopheles* mosquitoes for malaria control.

Translating either population replacement or suppression to *Anopheles* mosquitoes is not straightforward but mathematical models have already been developed to provide qualitative insights for designing optimal *Wolbachia* release strategies^7,8^. The *w*AlbB strain was transferred from *Ae. albopictus* into *An. stephensi* and resulted in significant *Plasmodium falciparum* inhibition and CI induction^9^, but associated mosquito fitness costs prevented further progression to field release trials. Recently the *w*Pip strain from *Culex quinquefasciatus* mosquitoes has been stably introduced into *An. stephensi* and despite high density infections there was minimal impact on mosquito fitness^10^ giving hope that this strain may provide more optimal phenotypic effects for release strategies. Given the technically challenging method of embryo microinjection to establish stable mosquito lines, adaptation of *Drosophila Wolbachia* strains to *Aedes* cell lines or using strains from closely related mosquito species were key reasons for successful *Ae. aegypti* transinfection^11^. Optimal candidate strains for species in the *Anopheles gambiae* complex or *Anopheles funestus* would be from within the *Anopheles* genera but historically *Wolbachia* was thought to not occur naturally within wild populations. We have recently discovered strains in genuine endosymbiosis in *An. moucheti* (*w*AnM) and *An. demeilloni* (*w*AnD) which were visualised in mosquito ovaries^12^. Our analyses also highlights that there is no evidence for *Wolbachia* in genuine symbiosis within major malaria vectors. Despite *Wolbachia* strains in the *An. gambiae* complex being at the threshold of detection, *Wolbachia* in *An. gambiae* and *An. coluzzii* have been shown to significantly reduce *Plasmodium* in naturally occurring populations^13,14^.

For *Wolbachia* to be used as a successful control strategy, strains must be identified, characterised for phenotypic effects and monitored in natural *Anopheles* populations from malaria-endemic countries. Currently surveillance detection methods used for *Wolbachia* include PCR amplification of *Wolbachia* specific genes or visualisation such as fluorescent *in situ* hybridisation. However, these are often expensive, time consuming and unless previous work has demonstrated evidence of *Wolbachia* strains in *Anopheles* populations, can result in insignificant evidence of a genuine endosymbiosis. Therefore, sequencing methods have been used to provide stronger evidence for the presence of genuine high-density *Wolbachia* strains in *Anopheles* mosquito populations^12^. Illumina or sanger sequencing have limited use due to high cost, the need for significant infrastructure and expertise in bioinformatics. Given these limitations, we explored the use of Oxford Nanopore Technology (ONT) as a viable tool to identify natural *Wolbachia* strains in *Anopheles* populations given this sequencing method is rapid and can be undertaken with minimal equipment opening the possibility of sequencing in malaria-endemic field settings. Marketed as fully portable, the MinION sequencer offers the possibility of being deployed in less-developed laboratories across the WHO African region, empowering local facilities with advanced capabilities for research and the ability to perform ‘in house’ sequencing negating the need to send samples to other research institutes.

Our study evaluated the usability and accuracy of ONT using the MinION for *Wolbachia* detection in *Anopheles* mosquitoes and assessed the additional equipment and facilities necessary to enable sequencing in field settings. Using more economical Flongle flow cells, we compared ONT sequencing using the portable Mk1b device across multiple *Anopheles* species comparing two library preparation kits: the Rapid PCR Barcoding Kit 24 V14 and 16S Barcoding Kit 24 V14. Our results showed consistent detection of *Wolbachia* within the two stably infected species (*An. demeilloni and An. moucheti)* but no evidence of *Wolbachia* strains in *An. gambiae*. Our results also showed higher average read counts for *Wolbachia* using the 16S Barcoding Kit 24 V14 with average read counts of 623 for *An. demeilloni* and 3440 in *An. moucheti*.

## Methods

### Sample collection and storage

Adult *Anopheles* mosquitoes of various species were collected from the DRC, Kenya and Cameroon with location and year of collection shown in **Table 1**. In Cameroon, adult female *Anopheles moucheti* females were collected in February 2024 using human landing catches^12^ from Olama Village (3.4125, 11.2841) and transported to OCEAC (Yaoundé) at 4°C for morphological identification. In Kenya, *Anopheles demeilloni* samples were collected in 2019 from Bigege Village (−0.5951053, 34.7488466) in Kisii, Western Kenya using CDC light traps and transported to KEMRI (Kisumu) for morphological identification. In DRC, *Anopheles demeilloni* samples were collected during ongoing entomological surveys in 2019 and 2024 using manual aspiration from Lwiro (−2.244097, 28.815232) and Maziba (−2.241816, 28.800961) in the Sud Kivu region of eastern DRC. Morphologically identified adult females from all collection locations were placed individually in 1.5 mL Eppendorf tubes, transported to the University of Warwick in temperature-controlled conditions and stored at -70°C prior to DNA extraction. Surface sterilisation using 70% ethanol and sterilised water washes was undertaken for all samples prior to DNA extraction.

**Table 1.**
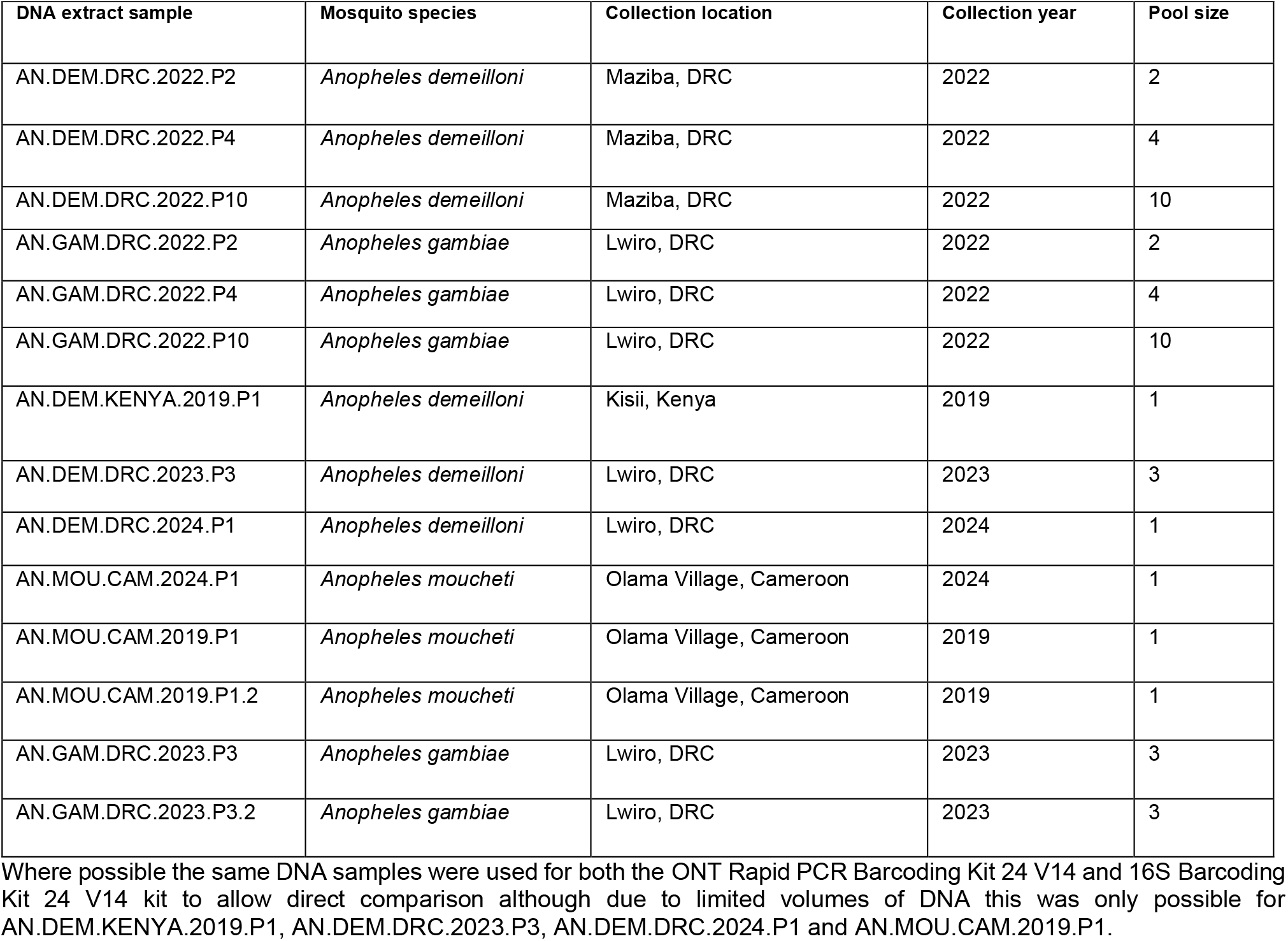
Species, collection information and pool size of *Anopheles* mosquito genomic DNA used for Library preparation using the ONT Field sequencing kit, ONT Rapid PCR Barcoding Kit 24 V14 and 16S Barcoding Kit 24 V14. AN= *Anopheles*, DEM= *demeilloni*, GAM = *gambiae*, MOU= *moucheti*, P= pool size.

| DNA extract sample | Mosquito species | Collection location | Collection year | Pool size |
| --- | --- | --- | --- | --- |
| AN.DEM.DRC.2022.P2 | *Anopheles demeilloni* | Maziba, DRC | 2022 | 2 |
| AN.DEM.DRC.2022.P4 | *Anopheles demeilloni* | Maziba, DRC | 2022 | 4 |
| AN.DEM.DRC.2022.P10 | *Anopheles demeilloni* | Maziba, DRC | 2022 | 10 |
| AN.GAM.DRC.2022.P2 | *Anopheles gambiae* | Lwiro, DRC | 2022 | 2 |
| AN.GAM.DRC.2022.P4 | *Anopheles gambiae* | Lwiro, DRC | 2022 | 4 |
| AN.GAM.DRC.2022.P10 | *Anopheles gambiae* | Lwiro, DRC | 2022 | 10 |
| AN.DEM.KENYA.2019.P1 | *Anopheles demeilloni* | Kisii, Kenya | 2019 | 1 |
| AN.DEM.DRC.2023.P3 | *Anopheles demeilloni* | Lwiro, DRC | 2023 | 3 |
| AN.DEM.DRC.2024.P1 | *Anopheles demeilloni* | Lwiro, DRC | 2024 | 1 |
| AN.MOU.CAM.2024.P1 | *Anopheles moucheti* | Olama Village, Cameroon | 2024 | 1 |
| AN.MOU.CAM.2019.P1 | *Anopheles moucheti* | Olama Village, Cameroon | 2019 | 1 |
| AN.MOU.CAM.2019.P1.2 | *Anopheles moucheti* | Olama Village, Cameroon | 2019 | 1 |
| AN.GAM.DRC.2023.P3 | *Anopheles gambiae* | Lwiro, DRC | 2023 | 3 |
| AN.GAM.DRC.2023.P3.2 | *Anopheles gambiae* | Lwiro, DRC | 2023 | 3 |

Where possible the same DNA samples were used for both the ONT Rapid PCR Barcoding Kit 24 V14 and 16S Barcoding Kit 24 V14 kit to allow direct comparison although due to limited volumes of DNA this was only possible for AN.DEM.KENYA.2019.P1, AN.DEM.DRC.2023.P3, AN.DEM.DRC.2024.P1 and AN.MOU.CAM.2019.P1.

### DNA extraction and quantification

Genomic DNA (gDNA) was extracted from adult female *Anopheles* mosquitoes using Qiagen DNeasy Blood and tissue kits according to manufacturer’s protocols. Surface Homogenisation of whole mosquitoes prior to DNA extraction was achieved using Qiagen Tissue lyser II (Qiagen, The Netherlands) at a speed of 25Hz for 5 minutes. Samples were used for library preparation as either individuals or in pools of three mosquitoes post-extraction if insufficient DNA concentration was obtained for individual samples (**Table 1)**. Before beginning ONT library preparation and standardising the concentration of DNA, DNA extracts were quantified using a Qubit 4 fluorometer (Invitrogen, Thermo-fisher scientific, Massachusetts, USA). For this 2μl of DNA extracts were individually combine with 198μL of the 1X Invitrogen dsDNA high sensitivity assay (cat no. Q33231) and placed in the Qubit 4 fluorometer to determine the concentration of DNA. This process was used for all DNA quantification stages throughout the study including post PCR quantification.

### ONT Field sequencing kit (SKU: SQK-LRK001) library preparation

DNA extracted from pools of mosquitoes (2, 4, 10 and 12 mosquitoes/pool) were analysed for total DNA to determine suitability for the Field sequencing kit (**Table 1**). Mosquito pool sizes of >10 only were considered to have enough genomic DNA (gDNA) to proceed with library preparation (400 ng of gDNA in 10 μL). In brief, 10 μL of gDNA was added to the Fragmentation Mix and then mixed, incubated at room temperature for 1 min and then at 80°C for 1 min. 10 μL of tagmented DNA was then added to the rapid adapter and mixed before incubated at room temperature for 5 min then immediately loaded onto the ONT flow cell.

### ONT Rapid PCR Barcoding Kit 24 V14 (SQK-RPB114.24) library preparation

Individual mosquito gDNA was prepared by diluting 1-5ng of DNA in nuclease free water to a volume of 3μL followed by combining with the fragmentation mix (FRM) and placed in a thermocycler (T100 Thermocycler, BioRad, California, USA) at 30°C for 2 min and then 80°C for 2 min. PCR was undertaken by combining the FRM/gDNA with NEB LongAmp Taq 2X master mix (cat no. M0287S) and each sample was combined with a different barcode (RLB) and underwent PCR amplification in a Bio-Rad T100 Thermocycler (California, USA) with initial denaturation (95°C for 3 min), followed by 25 cycles of 95°C for 15 sec, 55°C for 15 sec, and 65°C for 6 min with a final extension step of 65°C for 6 min. 25 cycles was determined by several preliminary trial PCRs as the protocol recommended 14 cycles produced too small a quantity of DNA to continue to library preparation. After PCR, EDTA was added to the reaction and DNA quantification was undertaken using an Invitrogen Qubit™ 1X dsDNA High Sensitivity (HS) assay (cat no. Q33232) and an Invitrogen Qubit 4 fluorometer. Each barcoded sample was then pooled in an equimolar ratio and combined with AMPure XP beads (AXP) at a 0.6x volume ratio. Samples were incubated on a hula mixer (TR-200 Rotator, Cole Parmer, Illinois, USA) and then pelleted on a magnetic rack (Ambion Magnetic Stand-96 AM10027, Invitrogen, Thermo-fisher scientific, Massachusetts, USA) and supernatant removed and discarded. Pellets were then washed twice with 80% ethanol solution and resuspended in elution buffer (EB) and then placed back on the magnetic rack where the eluate was retained. After the second quantification with the Invitrogen Qubit 4 fluorometer, 5-25fmol of sample was combined with EB to make up 5.5μL and combined 0.5μL of diluted rapid adapter (dRA). DNA libraries were then combined with sequencing buffer (SB) and library beads (LB).

### 16S Barcoding Kit 24 V14 (SQK-16S114.24) library preparation

Extracted gDNA from individual mosquitoes was prepared by diluting 10ng of DNA in 15μL of nuclease free water and then the diluted DNA was combined with NEB LongAmp Taq 2x master mix and respective 16S barcodes. Samples were submitted to PCR amplification in a Bio-Rad T100 Thermocycler (California, USA) with initial denaturation (95°C for 1 min), followed by 25 cycles of 95°C for 20 sec, 55°C for 30 sec, and 65°C for 2 mins with a final extension step of 65°C for 5 min. A volume of 4μL of EDTA was added to each barcoded sample before pooling and mixing with AMPure XP beads in 0.6x ratio. Samples were incubated on a hula mixer (TR-200 Rotator, Cole Parmer, Illinois, USA) and then pelleted on a magnetic rack (Ambion Magnetic Stand-96 AM10027, Invitrogen, Thermo-fisher scientific, Massachusetts, USA) and supernatant removed and discarded. The pellets were then washed twice with 80% ethanol solution and resuspended in elution buffer (EB) and then placed back on the magnetic rack where the eluate was retained. After the second quantification with the Qubit 4 fluorometer, 5-25fmol of sample was combined with EB to make up 5.5μL and combined 0.5μL of diluted rapid adapter (dRA). DNA libraries were then combined with sequencing buffer (SB) and library beads (LB).

### ONT sequencing

The Oxford Nanopore Technologies MinION Mk1C (a compact, portable self-contained device that combines the hardware for running nanopore sequencing experiments with fully integrated compute used for base calling and onward analysis) was first tested using pooled DNA libraries produced using the field sequencing kit (SKU: SQK-LRK001) on MinION flow cells (FLO-MIN004RA). A single flow cell run with a library prepared using the Rapid PCR Barcoding Kit 24 V14 (SQK-RPB114.24) was tested on the Mk1C but failed. As the Mk1C has been discontinued, all further ONT sequencing was performed using Flongle flow cells (FLO-FLG114) with a MinION Mk1B and flongle adapter powered by a Dell standard AIO desktop (210-BFWY) computer with an operating system >16 GB ram, a USB3 port and an internal or external storage of 1TB. Individual singular samples from gDNA extracted from an individual *Anopheles* mosquito were first tested and gDNA sample numbers increase on a single Flongle flow cell once consistent sequencing outputs were obtained. This was executed by using the barcoding features of each kit, barcoding different DNA samples before entering a final pool for sequencing. This additionally provided quality control testing of accuracy when barcoding multiple replicates of the same gDNA sample.

### Sequencing analysis

Analysis of the raw sequencing data output was achieved using the EPI2ME agent and portal software application. Within the EPI2ME agent, the MinKNOW software automatically saved all output files including FASTQ files that contain the sequence of nucleotide bases to be analysed. After sequencing completion, an output of FASTQ files was sorted by MinKNOW into FASTQ pass and FASTQ fail and passed files inputted into the EPI2ME agent analysis platform and assessed using a What’s In My Pot (WIMP) analysis for each individual run and split by barcode. Analysis was undertaken using the Centrifuge version 1.0.3-beta classification method, then results were filtered and combined to determine the counts of reads at species-level taxonomic assignments. For reads without a reliable species-level assignment, these were placed at ranks higher in the taxonomic trees but if reads were deemed unreliable, they were not included and reported as unclassified. A ‘Fastq 16S’ Bacterial analysis was employed for all runs using the 16S Barcoding Kit 24 V14 (SQK-16S114.24) but all failed with no reports available. Finally, phylogenetic analysis and tree construction was undertaken from WIMP analysis within the EPI2ME platform. Generated FASTQ files were also analysed using a SituSeq pipeline^15^ employing *DADA2*^16^. Following this pipeline, primers were trimmed from all reads, reads were then filtered and trimmed based on observed quality before being taxonomically assigned using Silva (Version 138.1 SSU)^17^ and Relative abundance explored at the genus level via bar charts created in *ggplot2*^18^.

## Results

### Total sequencing data output comparison of library preparation kits

Analysis of pooled mosquitoes (n=10) revealed total reads for *An. demeilloni* (sample AN.DEM.DRC.2022.P10) (**Figure 1A**) was 139,950 compared to 72,659 for *An. gambiae* (sample AN.GAM.DRC.2022.P10) (**Figure 1B**) when using the ONT Field sequencing kit libraries sequenced on the MinION Mk1C. The percentage of reads that were classified was similar (46.1%, 42.7% respectively) and of those that were classified both samples showed bacterial reads at greater than 60%. The requirement to use pooled mosquitoes (>10) to obtain sufficient gDNA, the cost of MinION flow cells and the discontinuation of both the field sequencing kit for library preparation and the MinION Mk1C devise prevented a more comprehensive analysis as to whether this could be applicable in malaria-endemic field settings. To assess the optimal method of ONT sequencing, two library preparation kits were compared: Rapid PCR Barcoding Kit 24 V14 and 16S Barcoding Kit 24 V14, using the more economical flongle flow cells (FLO-FLG114) and a Flongle adapter on a MinION Mk1B device on gDNA extracted from three *Anopheles* mosquito species: *An. gambiae, An. demeilloni* and *An. moucheti*. A total of 10/11 (91%) sequencing runs were successful with only a single run failing to produce any reads above the quality score threshold of nine (**supplementary tables S1-S2; Table 2**). The Rapid PCR Barcoding Kit library preparation allows sequencing of the whole genome in contrast to the 16S Barcoding Kit library preparation kit which only targets the ubiquitous bacterial *16S rRNA* gene. Overall, the Rapid PCR Barcoding Kit obtained a higher mean volume of data with 4/5 (80%) runs achieving over the theoretical max output of 2.6 GB for a Flongle flow cell (this was only achieved on one sequencing run for the 16S Barcoding Kit) (**Figure 2A**). On average the Rapid PCR Barcoding Kit generated 5.9Gb of data compared to only 1.5 Gb of data for the 16S Barcoding Kit. A similar trend was observed for the total amount of reads produced with 148,034 for the Rapid PCR Barcoding Kit and 63,854 reads for the 16S Barcoding Kit (**Figure 2B**). However, the most successful run (no.7) in terms of read number and data did result from the 16S Barcoding Kit 24 V14 producing 254,340 reads.

**Table 2.** ONT sequencing runs using flongle flow cells (FLO-FLG114) Samples tested were individually barcoded unless indicated and in some flow cell runs (eg. run 2) the same DNA sample was used in two individual barcodes and run in replicate on the same flow cell. Asterisk indicates run failure.

| Flow cell | Library preparation kit | Device | species | Samples tested | Barcode | Total reads | Q score | *Wolbachia* reads |
| --- | --- | --- | --- | --- | --- | --- | --- | --- |
| 1 | Rapid PCR | Mk1C | *An. demeilloni* | AN.DEM.DRC.2023.P3 | RLB1 | 400000 | 6.16 | 39 |
| 2 | Rapid PCR | Mk1B | *An. moucheti* | AN.MOU.CAM.2024.P1 | RLB2 | 6000 | 13.47 | 110 |
|  |  |  |  | AN.MOU.CAM.2024.P1 | RLB4 | 5000 | 13.42 | 110 |
| 3 | Rapid PCR | Mk1B | *An. demeilloni* | AN.DEM.KENYA.2019.P1 | RLB5 | 5669 | 12.71 | 0 |
|  |  |  |  | AN.DEM.KENYA.2019.P1 | RLB7 | 6701 | 12.54 | 2 |
| 4 | Rapid PCR | Mk1B | *An. moucheti* | AN.MOU.CAM.2019.P1 | RLB12 | 5000 | 13.1 | 29 |
|  |  |  |  | AN.MOU.CAM.2019.P1.2 | RLB18 | 13000 | 12.81 | 60 |
| 5 | Rapid PCR | Mk1B | *An. demeilloni* | AN.DEM.DRC.2023.P3 | RLB5 | 2905 | 13.25 | 1 |
|  |  |  |  | AN.DEM.DRC.2023.P3 | RLB6 | 10441 | 13.25 | 26 |
|  |  |  |  | AN.DEM.DRC.2023.P3 | RLB7 | 3985 | 13.25 | 7 |
| 6 | Rapid PCR | Mk1B | *An. gambiae* | AN.GAM.DRC.2023.P3 | RLB12 | 12393 | 12.03 | 0 |
| 7 | 16S | Mk1B | *An. moucheti* | AN.MOU.CAM.2024.P1 | RLB1 | 7788 | 11.49 | 1189 |
|  |  |  |  | AN.MOU.CAM.2024.P1 | RLB2 | 5901 | 11.49 | 757 |
|  |  |  |  | AN.MOU.CAM.2024.P1 | RLB3 | 7972 | 11.49 | 1278 |
|  |  |  |  | AN.MOU.CAM.2024.P1 | RLB4 | 10008 | 11.49 | 2809 |
| 8 | 16S | Mk1B | *An. moucheti* | AN.MOU.CAM.2019.P1 | RLB13 | 9709 | 12.4 | 5289 |
|  |  |  |  | AN.MOU.CAM.2019.P1 | RLB15 | 17689 | 12.4 | 14111 |
|  |  |  |  | AN.MOU.CAM.2019.P1.2 | RLB9 | 1649 | 12.4 | 1018 |
|  |  |  |  | AN.MOU.CAM.2019.P1.2 | RLB 12 | 1831 | 12.4 | 1070 |
| 9 | 16S | Mk1B | *An. demeilloni* | AN.DEM.DRC.2023.P3 | All RLBs | 67 | <9 | 30 |
| 10* | 16S | Mk1B | *An. demeilloni* | AN.DEM.DRC.2024.P1 | All RLBs | 0 | 0 | 0 |
| 11 | 16S | Mk1B | *An. demeilloni* | AN.DEM.DRC.2024.P1 | RLB1, RLB2 | 2183 | 10.25 | 1246 |
| 12 | 16S | Mk1B | *An. gambiae* | AN.GAM.DRC.2023.P3.2 | All RLBs | 4491 | 12.62 | 0 |

**Figure 1.**
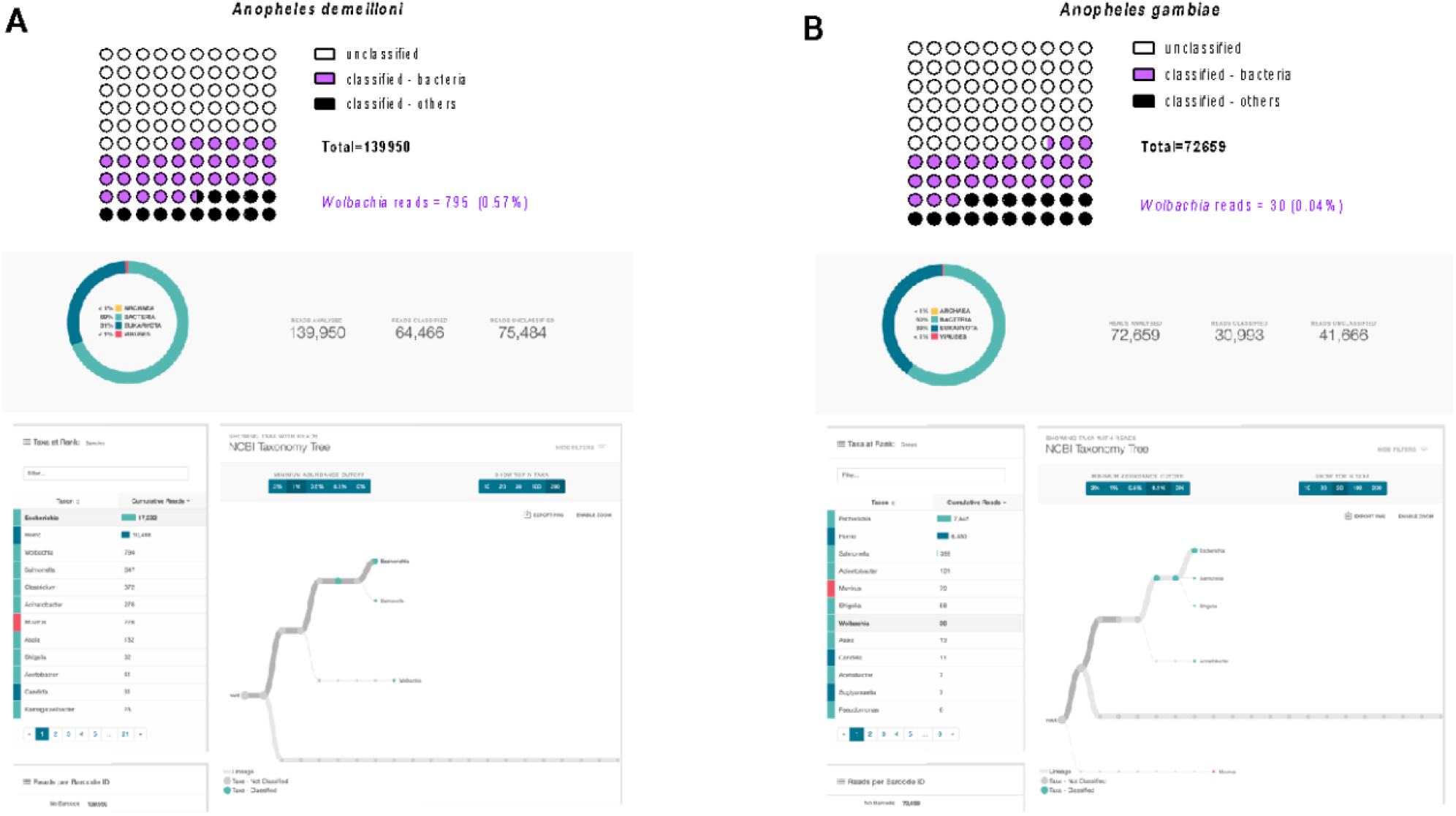
WIMP analysis of ONT sequencing using the field sequencing kit library preparations (n=10 mosquitoes/pool) from MinION sequencing on the Mk1C device. A) *Anopheles demeilloni* sample = AN.DEM.DRC.2022.P10, B). *Anopheles gambiae* sample = AN.GAM.DRC.2022.P10.

**Figure 2.**
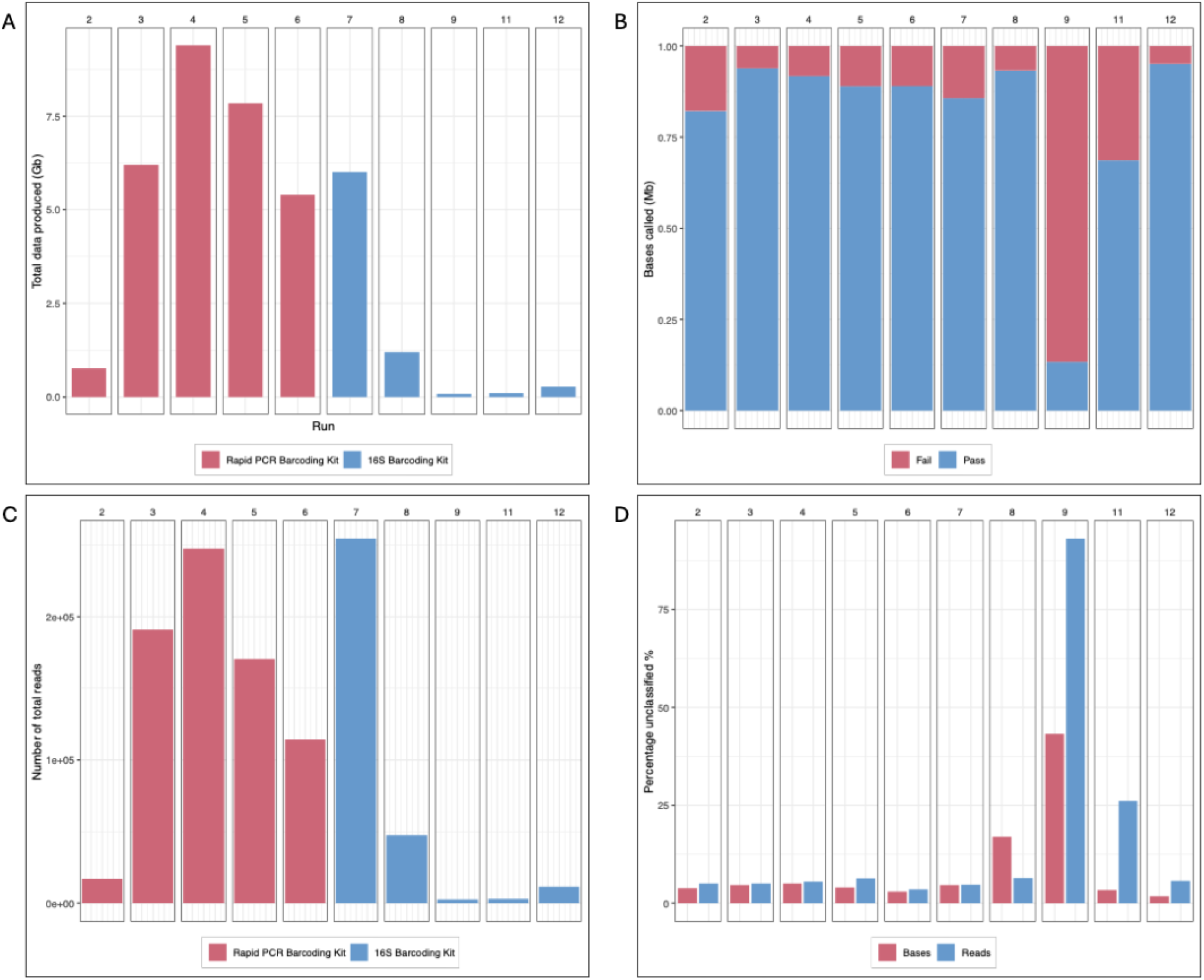
Evaluation of overall output ONT sequencing data for flongle flow cell runs on *Anopheles* gDNA samples. All bar graphs depict sequencing runs using two different barcoding kits: the Rapid PCR Barcoding Kit 24 V14 (runs 2-6) and the 16S Barcoding Kit 24 V14 (runs 7,8,9,11,12). Each run is represented by a vertical column. **A**) Total Data generated (GB). **B**) Percentage of passed vs failed bases called for each run. **C**) Total number of reads obtained in each sequencing run. **D**) Percentage of unclassified passed reads and unclassified passed bases for each run.

**Figure 3.**
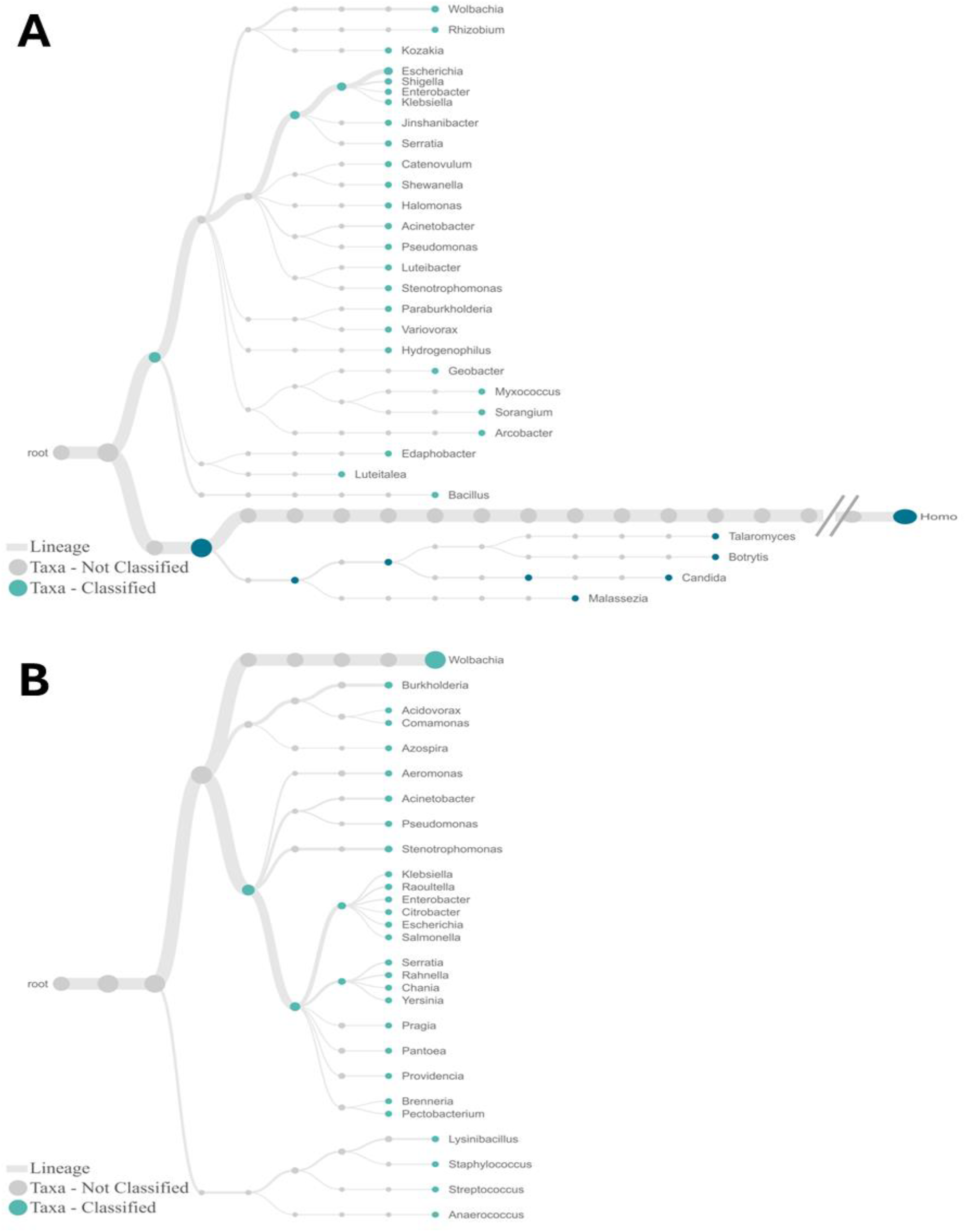
Phylogenetic Trees Illustrating Taxonomic Classification of Microbial Communities from WIMP analysis of *An. moucheti* (sample AN.MOU.CAM.2019.P1.2) **(A)** Library prepared with the Rapid PCR Barcoding Kit 24 V14 **(B)** Library prepared with the 16S Barcoding Kit 24 V14. Both phylogenetic trees show the top 30 genera at the cut off abundance 0.1%.

### Base calling failure rates

The percentage of bases that were called successfully where a pass was considered as a minimum quality score of 9 was also assessed. On average 10.8% of base calling failed when using the Rapid PCR Barcoding Kit compared to 28.8% of base calling failed when using the 16S Barcoding Kit (**Figure 2C**). Flonge flow cell run nine had the poorest quality with 86% of bases called falling below the quality cut-off score of nine. Removing run nine (which may have resulted from a technical error), the percentage base call failure rate for the 16S Barcoding Kit library preparations was still higher at 14.3%. Both Rapid PCR Barcoding Kit and the 16S Barcoding Kit library preparation methods utilised barcoding of mosquito gDNA samples, in cases where only one sample was used per run multiple barcodes of the same sample were used as per the protocol. On average the Rapid PCR Barcoding Kit had 5.16% and 4.16% unclassified reads and bases respectively (**Figure 2D**). In comparison, the 16S Barcoding Kit produced higher proportions of unclassified reads and bases at 27.24% reads and 14.02% bases unclassified. This demonstrates that a higher percentage of the sequencing data is lost due to failures in barcoding when using the 16S Barcoding Kit. Overall, on average when assessing the volume of data acquisition, total read number, proportion of passed bases and the succusses of the barcoding classify in the data the Rapid PCR Barcoding Kit was judged a higher quality method of library preparation. However, the most successful run (run 7) was using the 16S Barcoding Kit. The maximum samples running simultaneously was estimated to be four using flongle flow cells as beyond this we were not able to produce a high enough read counts whereby results would be accepted using the quality cut off scores.

### Comparison of ONT sequencing

Sequencing data output from libraries generated from the field sequencing library kit (MinION flow cell - R9.4.1) and libraries generated from the Rapid PCR Barcoding Kit 24 V14 (Flongle flow cell -FLO-FLG114) with identical sequencing run lengths of 24 hours were compared. The MinION flow cell (R9.4.1) theoretically produces approximately 18.5 times the volume of data than the Flongle flow cell (FLO-FLG114). However, when compared to the results achieved from the Flongle flow cell runs (**Table 3**), runs 4,5, and 6 achieved greater volumes of data. However, this was consistent for all runs as sequencing data output showed significant variability from 0.1 to 9.39 Gb on average. In 4/5 (80%) of sequencing runs undertaken on libraries generated from the Rapid PCR Barcoding Kit and sequenced on the Flongle flow cell, more bases were produced than sequencing on the MinION flow cell. In contrast, libraries generated from the 16S Barcoding Kit only produced a greater number of bases than the MinION flow cell in one instance (run 7). Although sequencing on the MinION flow cell achieved 100% of reads called this was not achieved in any of the Flongle flow cell runs, with some as low as 8.74% (run 4) with the highest % reads called for a 24-hour run at 95.71%. The MinION flow cell also achieved a higher number of bases passed than any Flongle flow cell run and had a higher proportion of reads called and greater amount of passed called bases. However, sequencing on Flongle flow cells produce similar numbers of estimated bases, reads generated, and total data produced.

**Table 3.** ONT sequencing output comparison for run duration of 24 hours comparing three library preparation kits.

| Sample(s) | Flow cell | Library prep kit | Estimated bases (Mb) | Reads generated | Total data (Gb) | Reads called (%) | Bases passed (Mb) | Bases failed (Mb) |
| --- | --- | --- | --- | --- | --- | --- | --- | --- |
| AN.DEM.DRC.2022.P10<br>(*An. demeilloni*) | MinION<br>(R9.4.1) | Field | 290 | 251010 | 5.3 | 100 | 86.08 | 171.72 |
| AN.MOU.CAM.2024.P1<br>(*An. moucheti*)<br>AN.MOU.CAM.2019.P1<br>(*An. moucheti*) | Flongle<br>(FLO-FLG14) | Rapid<br>PCR | 699 | 247480 | 9.4 | 8.74 | 59.32 | 5.31 |
| AN.DEM.DRC.2023.P3 x2<br>(*An. demeilloni*) | Flongle<br>(FLO-FLG14) | Rapid<br>PCR | 613 | 170480 | 7.8 | 10.14 | 58.50 | 7.24 |
| AN.MOU.CAM.2024.P1 x4<br>(*An. moucheti*) | Flongle<br>(FLO-FLG14) | 16S | 374 | 254340 | 6.0 | 15.00 | 51.17 | 8.57 |
| AN.DEM.DRC.2023.P3 x2<br>(*An. demeilloni*) | Flongle<br>(FLO-FLG14) | 16S | 6 | 2630 | 0.1 | 95.71 | 0.17 | 1.11 |

### Overall microbiome relative abundance

Analysis using WIMP workflow from the EPI2ME platform was utilised for libraries sequenced using the 16S Barcoding Kit to determine the relative abundance of bacteria in *An. moucheti, An. demeilloni* and *An. gambiae*. Initially the consistency of the microbiome profile using four replicate samples of *An. moucheti* (sample AN.MOU.CAM.2024.P1) was assessed with consistent profiles generated for each barcode (**Figure 5A**). Across all replicate libraries, the bacterial community composition appeared consistent, with *Wolbachia* being the most dominant bacteria. *Enterobacter, Escherichia* and *Klebsiella* also were significant bacteria genera in this sample. Following a SituSeq pipeline, taxonomy was assigned to 16S rRNA reads using Silva database. The same consistency as described previously was observed between different barcodes for sample AN.MOU.CAM.2024.P1 (**Figure 5B)**. The number of reads pre- and post-quality filtering can be found in Table 4. All barcodes resulting in fewer than ten initial reads were removed from the dataset.

**Table 4.**
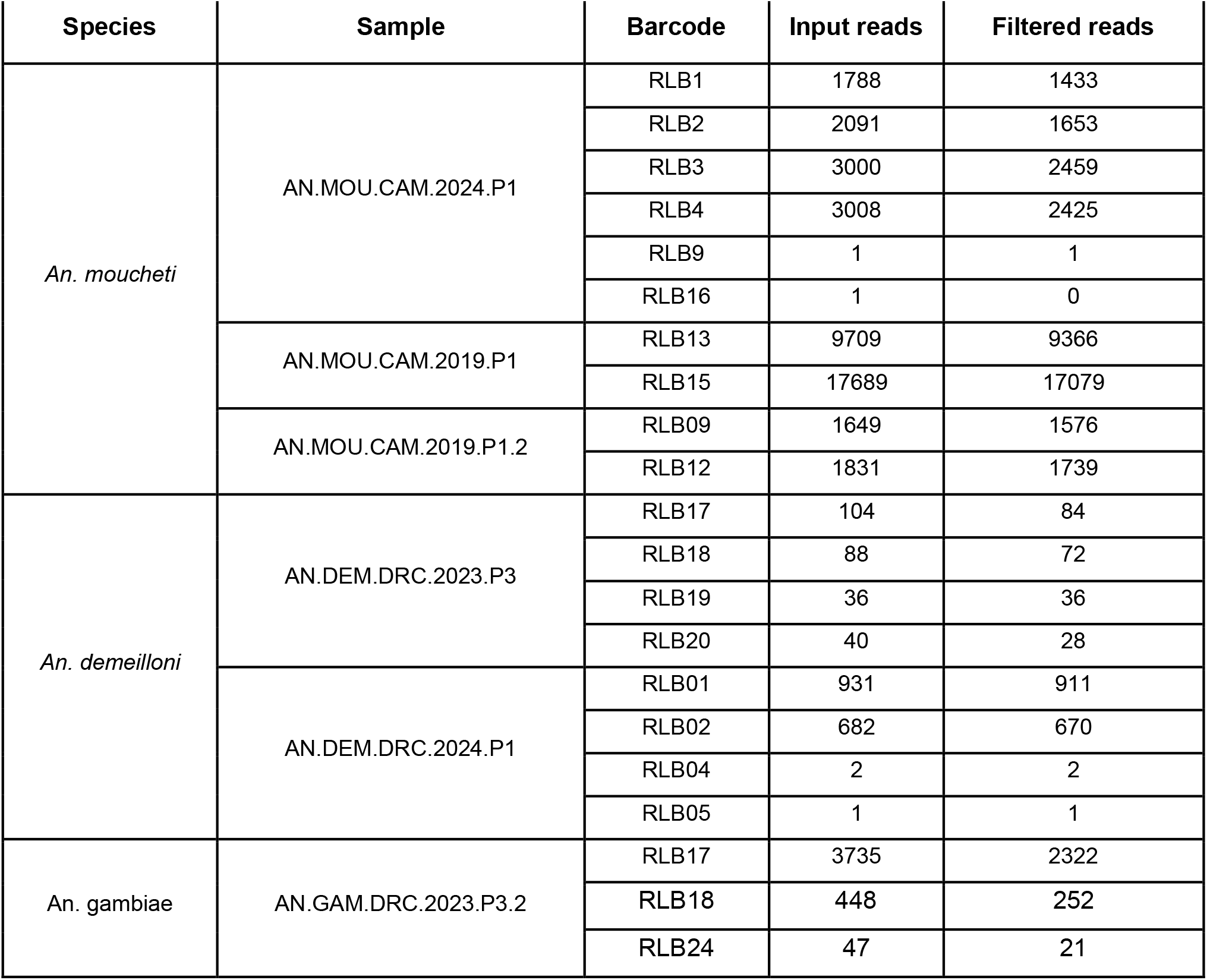
Number of reads generated per barcode using 16S rRNA nanopore sequencing kit for individual *Anopheles* mosquitoes. Input reads show the number of reads generated by ONT sequencing and filtered reads show those that passed SituSeq pipeline quality filters.

**Figure 4.**
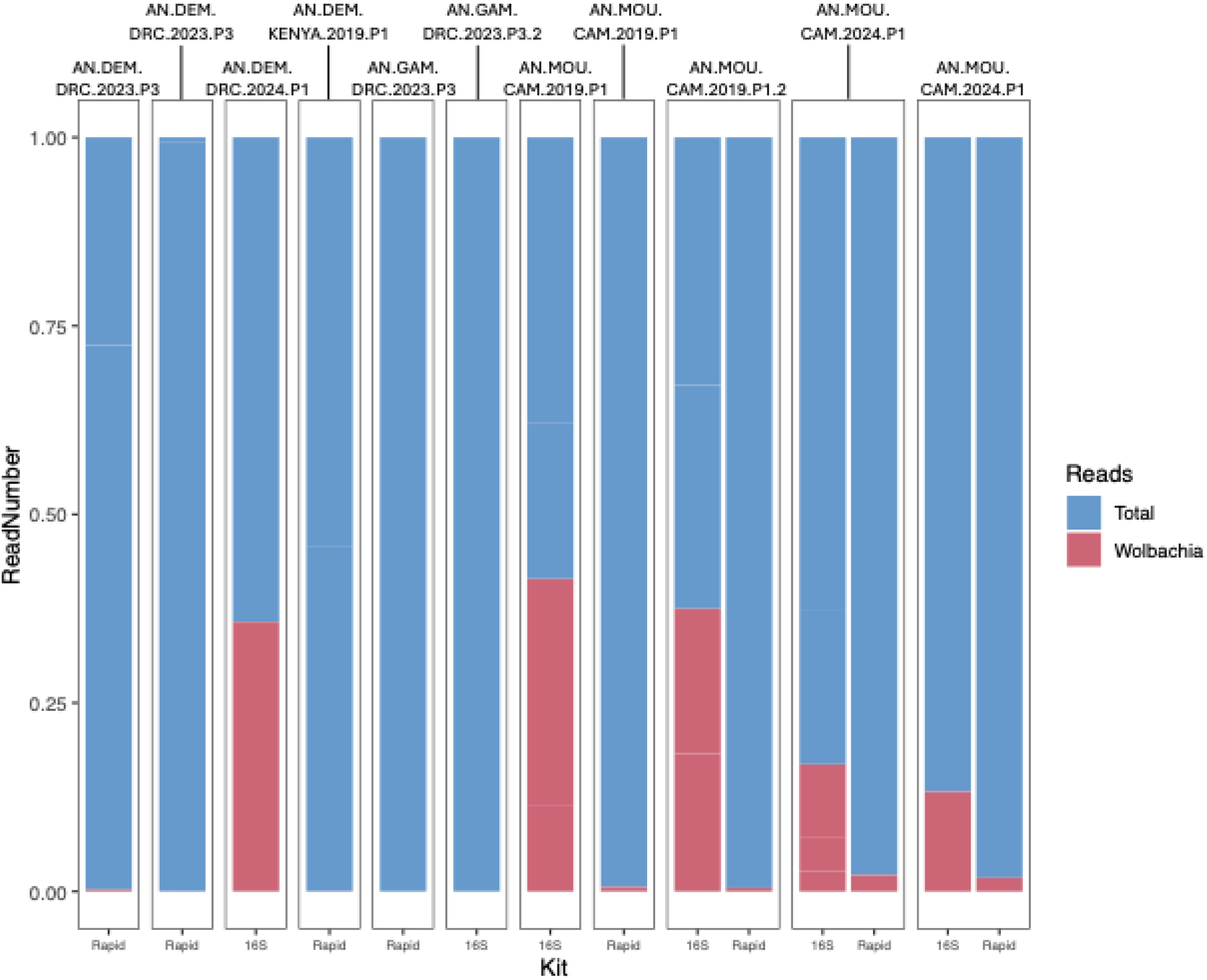
Proportion of total reads assigned to *Wolbachia* from ONT Flonge flow cell sequencing. Libraries prepared and sequenced using either the Rapid PCR Barcoding Kit 24 V14 (Rapid) or 16S Barcoding Kit 24 V14 (16S).

**Figure 5.**
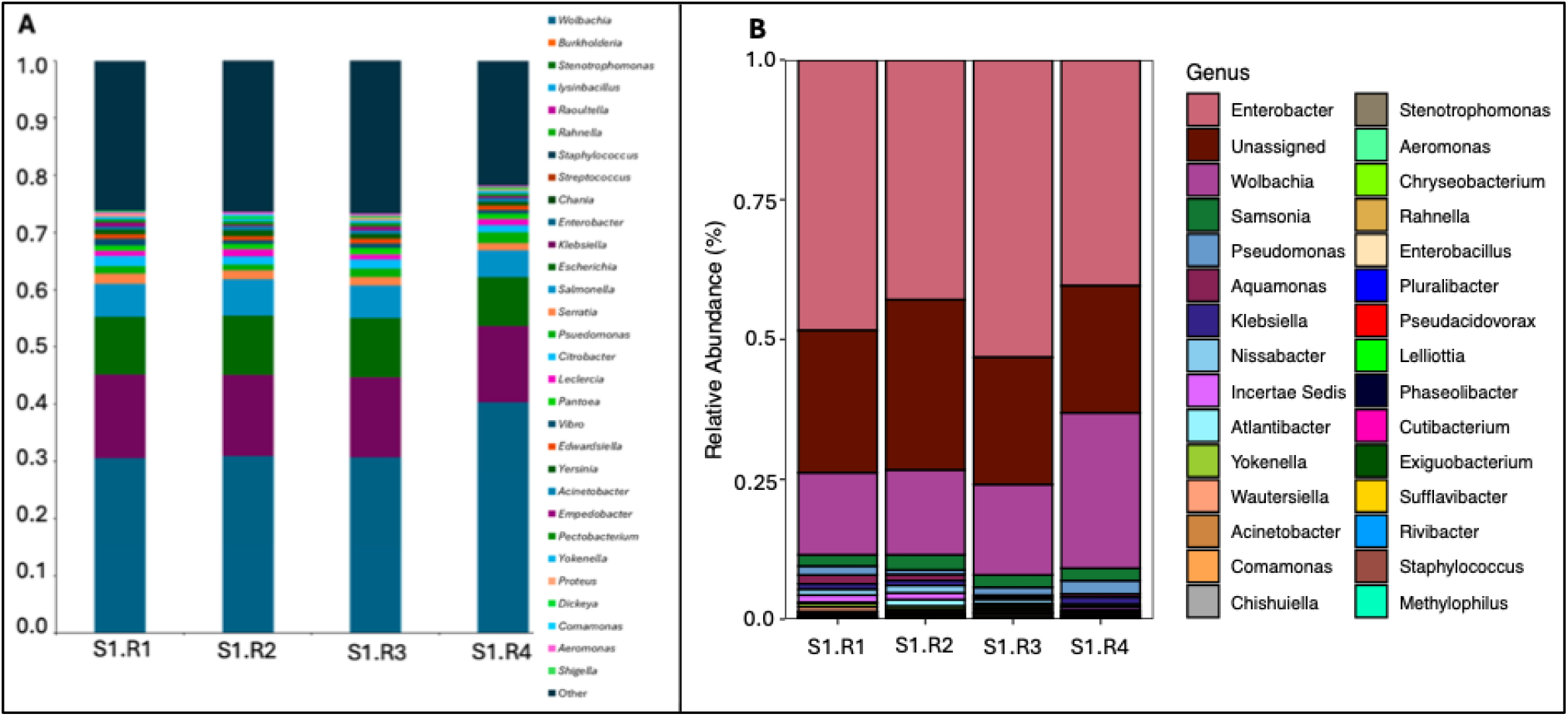
The microbial composition of sample AN.MOU.CAM.2024.P1 across four sequencing replicates using the 16S rRNA Barcoding Kit 24 V14. Output was analysed using two different pipelines: A) Epi2Me and B) SituSeq. Bacterial genera are listed in order of relative abundance. “Other” represents the sum relative abundance of bacterial genera that fell outside of the top 30 most abundant. “Unassigned” refers to all reads that were not assigned to the genus level following the SituSeq pipeline.

**Figure 6.**
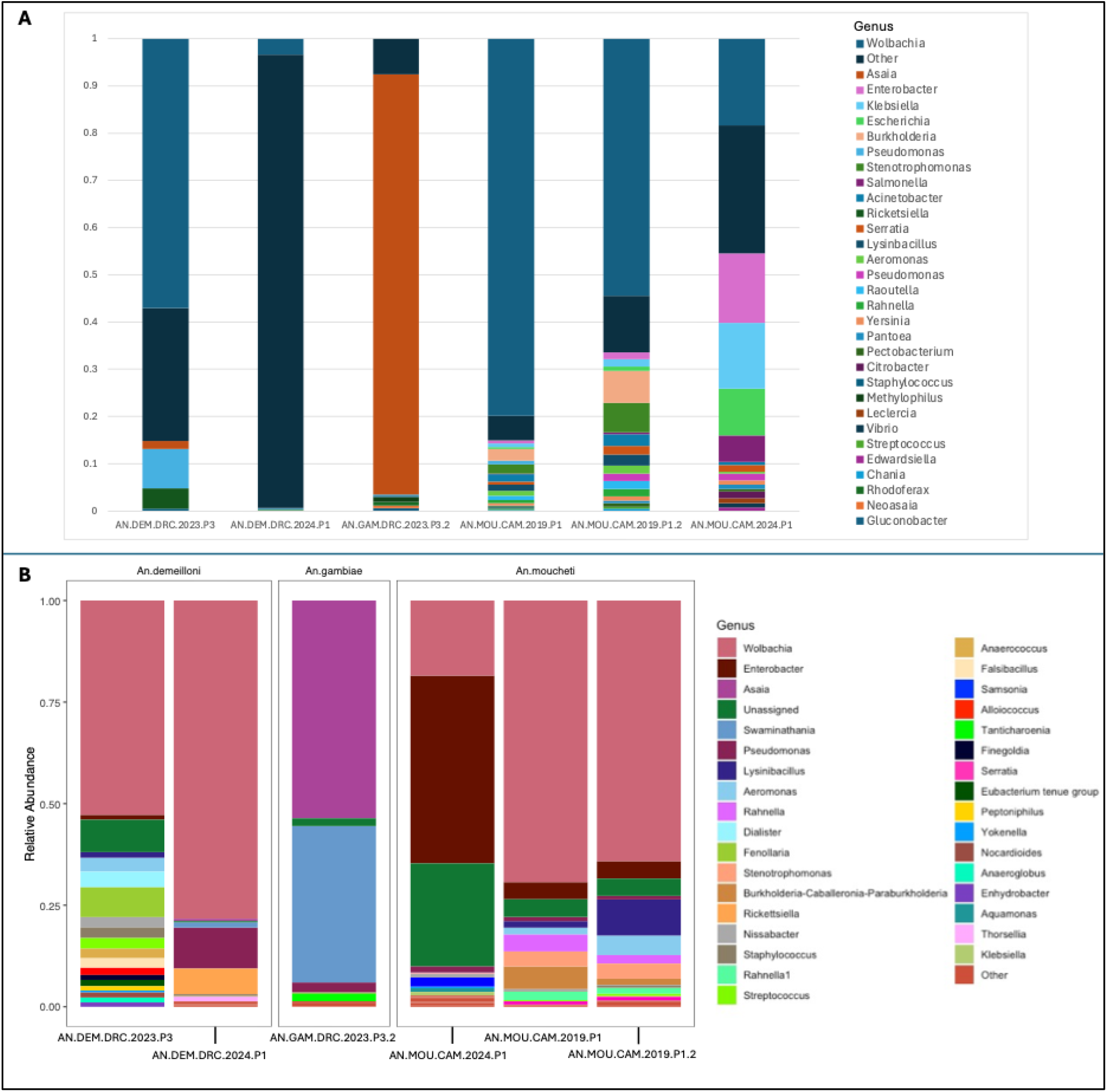
Relative abundance of bacterial genera within the *An. demeilloni, An. gambiae* and *An. moucheti* microbiome assessed using using the 16S rRNA Barcoding Kit 24 V14. Output was analysed using two different pipelines: A) Epi2Me and B) SituSeq. Bacterial genera are listed in order of relative abundance. For plot A, “Other” represents the sum relative abundance of bacterial genera that fell outside of the top 30 most abundant. In plot B, “Unassigned” refers to all reads that were not assigned to the genus level following the SituSeq pipeline and “Other” represents all bacterial genera that had a relative abundance of less than one percent of overall reads.

Taxonomic assignments were then assessed in terms of relative abundance (Figure 7) following the Epi2ME **(Fig 7A)** or SituSeq **(Fig 7B)** pipelines. There were several consistent findings between the two pipelines. *Wolbachia* dominated the *An. moucheti* and *An. demeilloni* microbiome and was not identified in the *An. gambiae* microbiome. Conversely, the genus *Asaia* had the highest relative abundance in *An. gambiae*, but was not identified in either of the other *Anopheles* species. There was a high relative abundance of Enterobacter in sample AN.MOU.CAM.2024.P1 observed in the output of both pipelines, with a slightly higher relative abundance estimated using the SituSeq pipeline. The single *An. gambiae* sample included (AN.GAM.DRC.2023.P3.2) was dominated by *Asaia* in both resulting analyses, however when assessing the SituSeq pipeline a high relative abundance of *Swaminathania* was also seen, which was not identified using Epi2ME analysis. When assessing the reads using the SituSeq pipeline, there was a relatively high proportion of unassigned reads in the majority of samples, with this group having the fourth highest overall relative abundance, however, this did vary between individual samples.

### *Wolbachia* detection using MinION flow cell sequencing

Using a library generated from the ONT Field sequencing kit sequenced on the MinION Mk1C, pooled *An. demeilloni* mosquitoes (sample AN.DEM.DRC.2022.P10) had total reads of 139,950 of which 69% were bacteria (**Figure 1**). The cumulative read count indicated the most abundant bacterial genera was *Escherichia* followed by *Wolbachia* (794 reads) which equated to 0.57% of total reads. Interestingly, the genus *Asaia* (known to compete with *Wolbachia* in *Anopheles* lab populations for colonisation of the germline ^19^, was present with 182 reads but as this library preparation kit required pooling 10 mosquitoes to obtain sufficient gDNA this does confirm co-infection with *An. demeilloni* individual mosquitoes. Pooled *An. gambiae* mosquitoes (sample AN.GAM.DRC.2022.P10) resulted in total reads of 72,659 with 60% identified as bacteria. In contrast to *An. demeilloni*, only 30 *Wolbachia* reads were identified in *An. gambiae* which equated to only 0.04% of total reads. *Wolbachia*- designated reads were ∼13 fold more abundant in *An. demeilloni than An. gambiae* which is consistent with other methods comparing *Wolbachia* densities in these two species such as quantitiave PCR and *16S rRNA* sequencing^12^.

### *Wolbachia* detection using flongle flow cell sequencing

The number of *Wolbachia* reads identified using WIMP analysis (**Figure 4**) from *An. demeilloni, An. moucheti* (known to also have a stable *Wolbachia* strain - *w*AnM) and *An. gambiae* were compared on the more economical flongle flow cells and a Flongle adapter on a MinION Mk1B device. For libraries prepared using the Rapid PCR Barcoding Kit only an average of 42 *Wolbachia* reads were detected for *An. demeilloni* and *An. moucheti* (**Table 2**). In contrast, libraries prepared using the 16S Barcoding Kit averaged significantly more with 2,038 reads. In terms of abundance (*Wolbachia*/total reads), the 16S Barcoding Kit average was 0.42 compared to only 0.006 for the Rapid PCR Barcoding Kit library preparation method. Neither library preparation method used in combination with flongle flow cells resulted in any *Wolbachia* reads in *An. gambiae*.

## Discussion

Overall, ONT sequencing using libraries prepared with the field sequencing kit and run on MinION flow cells resulted in a higher proportion of reads called and greater amount of passed called bases than the Flongle flow cell runs using either the Rapid PCR Barcoding Kit or 16S Barcoding Kit library preparation method. However, ONT sequencing on the Flongle flow cells were able to generate similar numbers of estimated bases, reads generated, and total data produced. Our results suggest that if the main data output desired is detection of *Wolbachia* in *Anopheles* using flongle flow cells then the 16S Barcoding Kit library preparation method is optimal given this averaged 2,038 *Wolbachia* reads across *Anopheles* species (*An. demeilloni* and *An. moucheti*) that have previously been shown to have *Wolbachia* strains in genuine symbiosis^12^. This was expected as the Rapid PCR Barcoding Kit libraries result in both the mosquito and microbiome rather than the 16S Barcoding Kit which targets just the bacterial microbiome.

The 16S Barcoding Kit library preparation method was also able to provide consistent microbiome profiles in which *Wolbachia* was the dominant species which is consistent with previous profiles using *16S rRNA* Illumina amplicon sequencing^12^. The absence of *Wolbachia* reads in *An. gambiae* from flongle flow cell ONT sequencing is also consistent with previous Illumina sequencing and PCR based studies not providing conclusive evidence for the presence of a genuine natural *Wolbachia* strain in this species^12,20^. However, *Asaia* was detected in *An. gambiae* (sample AN.GAM.DRC.2023.P3) with a relative abundance of 89% providing further evidence for a potential competitive mutual exclusion between these two bacteria. *Wolbachia* strains in *Anopheles* have been shown to be inversely correlated to *Asaia* which is stably associated with several *Anopheles* species^19, 21-24^. There are limited studies reporting co-infections in wild *Anopheles* populations^23^ but there were small numbers of individual co-infections from *An. melas* and *An. gambiae* s.s.–*melas* hybrids collected in Guinea^24^.

ONT sequencing has been used previously in field settings due to its portability, price, and ease of use for local/in-field sequencing. For example, in 2015 ONT was used to sequence samples that contained Ebola virus during the largest outbreak on record^25^. The use of ONT for vector-borne disease surveillance is increasing with ONT sequencing of *P. falciparum* malaria parasites from dried blood spots has been developed that can detect key drug-resistance associated genes, diagnostic test antigens and polymorphic markers^26-29^. Mosquito genome sequencing is also possible as ONT contributed to the development of an improved chromosome-scale genome assembly for *Culex quinquefasciatus* (the principal vector of West Nile virus)^30^ and produced de novo chromosome-level genome assemblies for *An. coluzzii* and *An. arabiensis*^31^. Mosquito bloodmeal analysis has been undertaken using ONT^32^ and ONT sequencing has also been tested for arbovirus surveillance^33^ including to identify yellow fever virus (YFV) in mosquitoes collected from Brazil during an outbreak in 2021^34^. Ross River virus screening was undertaken on *Aedes* mosquito RNA in 2017 but required the use of MinION flow cells and was less accurate compared to illumina amplicon sequencing due to a high error rate^35^. Large scale ONT sequencing (using flongle flow cells) on *Anopheles* mosquitoes analysed over 6,000 samples collected between 2003 and 2019 for the presence of Bagaza virus^36^.

For detecting *Wolbachia* strains in *Anopheles* mosquitoes, the 16S Barcoding Kit does have limitations for use in field settings such as the need for laboratory equipment for DNA extraction and library preparation. *Wolbachia* has recently been detected using ONT in fleas and ticks (*Rhipicephalus sanguineus* and *Ctenocephalides felis felis*)^37^ using the Ligation sequencing gDNA kit (SQK-LSK109). In addition, ONT sequencing from libraries prepared using the Ligation sequencing gDNA kit has been used to identify a *Wolbachia* bacteriophage (phage WO) of the southern ground cricket (*Allonemobius socius*)^38^ but unfortunately this kit has been discontinued limiting comparison to our study. Despite ONT having the advantage of requiring less infrastructure than other methods of sequencing, to generate libraries using the 16S barcoding kit the minimum equipment needed is a thermocycler, hula mixer, magnetic racks and vortex machines. The flongle flow cells run on the MinION Mk1B device also does require significant computing power, with a minimum storage requirement of 1TB, which may not be achievable in field settings. The barcoding kits used allowed for the testing of a theoretical maximum of 24 samples but due to limitations in read numbers we found only four *Anopheles* mosquito samples could be used per flongle flow cell run limiting sample numbers. ONT does has the advantage of variable read lengths and ultra-long reads (>1Mb) being permits whole genome sequencing. This could provide a rapid method of sequencing *Wolbachia* strain genomes that are present in wild *Anopheles* populations as to date only two strains have been sequenced^39^. ONT sequencing quality could be further improved by extraction of *Anopheles* DNA using kits that target high molecular weight DNA (something not testing in our study). Malaria disproportionately affects sub-Saharan Africa that is facing great economic challenges; therefore a more economically viable ‘in-field’ sequencing method may be crucial to novel vector control strategies such as those using *Wolbachia* bacteria.

## Supplementary information

**Table S1.**
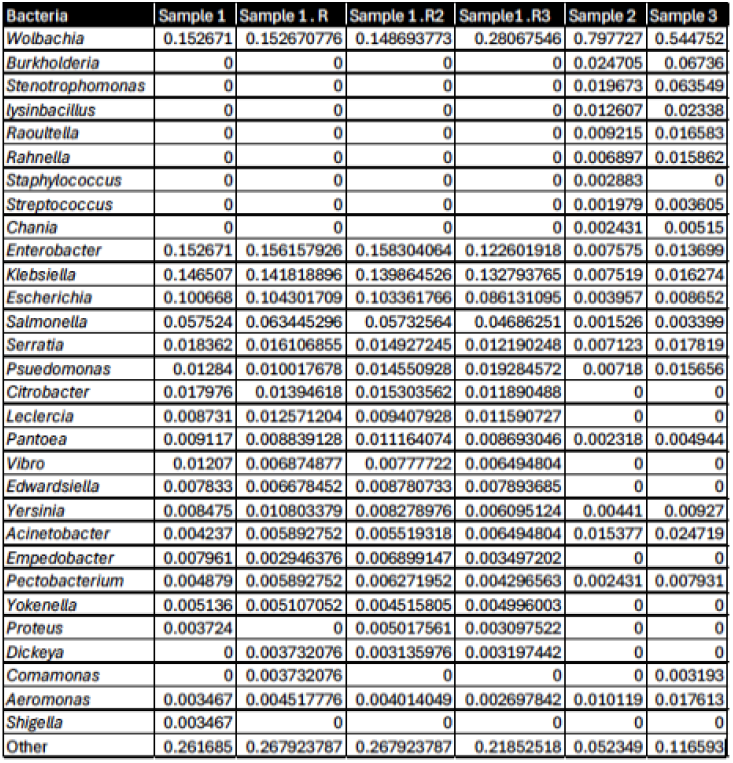
Relative abundance of bacteria from *An. moucheti* samples from ONT flongle flow cell sequencing using libraries prepared using the 16S barcoding kit. Sample 1 = AN.MOU.CAM.2024.P1, Sample 2 = AN.MOU.CAM.2019.P1, Sample 3 = AN.MOU.CAM.2019.P1.2.

**Table S2.**
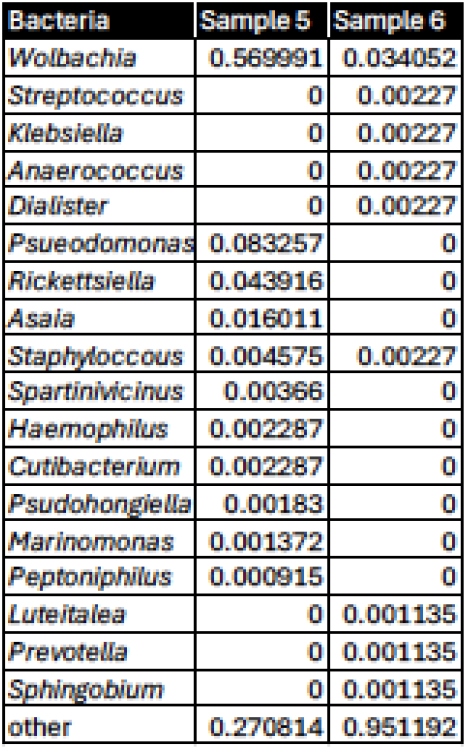
Relative abundance of bacteria from *An. demeilloni* samples from ONT flongle flow cell sequencing using libraries prepared using the 16S barcoding kit. Sample 5 = AN.DEM.DRC.2023.P3,, Sample 6 = AN.DEM.DRC.2024.P1.

**Table S3.**
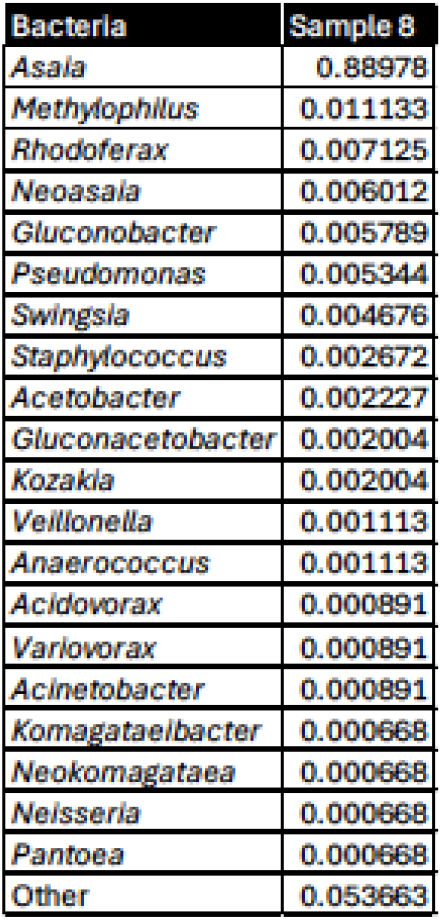
Relative abundance of bacteria from *An. gambiae* from ONT flongle flow cell sequencing using libraries prepared using the 16S barcoding kit. Sample 8 = AN.GAM.DRC.2023.P3.

